# High performance, large-scale multi-compartment Hodgkin-Huxley simulation of Drosophila’s whole-brain neural circuit model

**DOI:** 10.1101/2022.11.01.512969

**Authors:** Kaoruko Higuchi, Tomoki Kazawa, Buntaro Sakai, Shigehiro Namiki, Stephan Shuichi Haupt, Ryohei Kanzaki

**Author notes:** The first two authors contributed equally to this work.

## Abstract

A major challenge in neurosciences is the elucidation of neural mechanisms in brains that are crucial for the processing of sensory information and the generation of adaptive behavior. In conjunction with the ever-growing body of experimental data, computational simulations have become crucial in integrating information and testing hypotheses, requiring fast large-scale simulators. We constructed a whole-brain neural circuit model of the fly Drosophila with biophysically detailed multi-compartment Hodgkin-Huxley models based on the morphologies of individual neurons published in open databases. Performance tuning of the simulator enabled near real-time simulation of the resting state of the Drosophila whole-brain model in the large-scale computational environment of the supercomputer Fugaku, for which we achieved in excess of 630 TFLOPS using 480k cores. In our whole-brain model, neural circuit dynamics related to a standard insect learning paradigm, the association of taste rewards with odors could be simulated.

## I. Introduction

Animal brains control a wide variety of complex and adaptive functions such as integration of sensory information from multiple modalities, generation of adaptive behaviors in complex natural environments, learning and memory. However, it remains unclear which neural circuit properties and activity patterns in animal brains exactly control these functions. The elucidation of such principles is one of the most important issues in neuroscience. In recent years, computational simulations based on connectomes have become feasible as a bottom-up approach to understand the information processing in brains [1] [2] [3], since such models can integrate results obtained using a variety of different experimental approaches, which include the studies of single neuron physiology, the observation of neurons’ collective behaviors, and many histological methods. It is required to reconstruct, on a computer, neural circuits which are biophysically detailed models that replicate the physical world, and the multi-compartment Hodgkin-Huxley model [4] has become a typical approach used in the construction of such models [5].

In this study, we focused on insect brains, specifically that of Drosophila, to build a large-scale simulation using multi-compartment Hodgkin-Huxley models. The reason was as follows. First, the number of neurons in the insect brain is about 10^4^ to 10^6^, which is only one in a million compared to humans, but it shows characteristics of fairly good adaptability in complex environments executing path integration during search flights, possessing spatial memory [6], being capable of odor-taste association learning [7] and being able to generate numerous other sophisticated behaviors. Conceptually we could simulate such brain functions of the insect brain with less computational resources than in the case of the vertebrates. In addition, due to the smaller number of neurons, we considered it possible to perform detailed real-time simulations of the entire brain, given the capability of currently available supercomputers. Second, in an insect brain, the role and functions of a single neuron are more important than that of a mammalian counterpart. For insect brains, the existence of neurons with unique and complex morphology, that play an important roles in specific adaptative behaviors led to the “Identified neuron” doctrine and give an advantage for including each neurons’ morphology into the neural circuit simulation of insect brains [8] [9]. In some cases sub-regions of single neurons have been observed to have distinct roles in insect brains [10] [11] in addition to the dendric subcellar integration observed in vertebrate neurons [12]. Accordingly, it is crucial to incorporate the cells’ morphological information in the simulation of the insect brain. Drosophila is an insect whose neurons are fairly well studied, and morphological information at single neuron level is available in public databases.

Brains have interfaces as a sensor and a control-system to the real world that are too complex and dynamical to calculate easily. The real-time simulation is one of the targets used for the execution speed of computer simulation in the field of neuroscience. The significance of realizing real-time simulation speed is that it becomes possible to create a closed-loop experimental environment, in which the constructed brain model can be connected to a robot in a real environment and its behavior is controlled by the simulator. Also, real time data assimilation under the condition that the simulation processes information from the real outside world may become possible. To execute simulations using such detailed models at a large scale, it is indispensable that a simulator which can process information at high speed on modern supercomputer architectures with multi-level parallelism is made available. The development of such a neural simulation is also significant in that it can serve as a common platform for whole-brain simulations of various organisms.

## II. Related work

The first work of whole brain scale simulation was about the human thalamocortical system with 1 million neurons (1/10000 scale) using a combination of passive cable models and Izhikevich spike generator models that implemented the macroscopic pathway and microcircuit in cortex and thalamus [1]. This simulation exhibits oscillatory activity and its propagation as a steady state of natural brain. For the cortex, larger scale simulations of 10^9^ neurons using Integrate-and-fire models [13] model as our neuronal model. Other models commonly [14] or more detailed simulations of 30k neurons using multi-compartment Hodgkin-Huxley models for dendritic integration and spike generation [2] were executed in the modern HPC systems. These have been milestones towards the complete whole brain simulation in silico though we are still a long way from simulating whole brain functions at realistic speed to improve their understanding and for real-world applications. The performance aspect in massively parallelized environments has recently received some attention. NEURON K+ [15] using multi-compartment Hodgkin-Huxley models achieved 187 TFLOPS/200k cores on the K computer in a neural circuit simulation using (OpenMP + MPI) Hybrid Parallel, Structures-of-Array memory (SoA) layout of gating variables, and using MPI_send/receive for the implementation of synaptic transmissions. This has been a special case of neurons and neural circuits in an insect brain but projects to re-design neuron/neural circuit simulators for high performance in massively parallel environment have been initiated, such as Arbor [16] and CoreNEURON [17]. At least SoA memory layout introduced by both resulted in 5 times or more speed up in all cases in symmetrical clusters or hybrids between CPU and GPGPU.

The first study to simulate a Drosophila whole-brain model [3] was carried out at Tsing Hua University in Taiwan. The brain model consisted of 20,089 neurons extracted from the FlyCircuit database [18], and determination of the polarity of neurons and connectivity estimations were performed. The neurons were modeled using a leaky integrate-and-fire(LIF) model. The resting state simulation was conducted using an Intel CPU at 3.6 GHz (E3-1270v5), achieving 1/35 real-time speed. While the LIF model is computationally inexpensive, it is one of the simplest models that approximates a single cell as a cell body. In insects, where the spatiotemporal activity of a single neuron may play an important role, more detailed modeling may be required.

## III. Optimization of Neural simulator

### A. Model framework: multi-compartment Hodgkin-Huxley type model

We adopted the multi-compartment Hodgkin-Huxley used in neuroscience simulations are Leaky Integrate-and-Fire (LIF) model and Izhikevich model [19]. However, because the role of compartments of a single cell is vital in insects as mentioned in I, we have judged that more detailed modeling which includes morphological information is necessary for more accurate simulation. We selected the multi-compartment Hodgkin-Huxley model as it best suited our purpose, synaptic contacts were implemented using the double exponential decay synapse model.

In the multi-compartment Hodgkin-Huxley model, one cell is divided into multiple parts, where each part is approximated by cylindrical shapes called compartments. The model calculates the potential change in each compartment using the Hodgkin-Huxley equation, and calculates the potential propagation between compartments using the cable equation [20] [21](Fig.1).

**Fig. 1:**
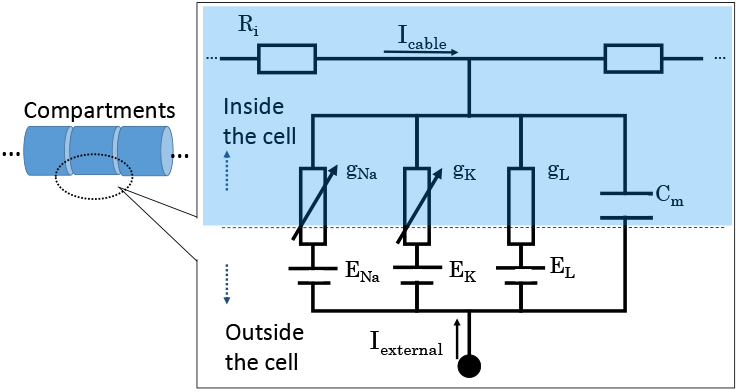
Schematic diagram of multi-compartment Hodgkin-Huxley model. Cytoplasm indicated in light blue.

The Hodgkin-Huxley model describes the dynamics of ion channels using differential equations and expresses the temporal changes in membrane potentials V using these equations. It consists of the following four equations.

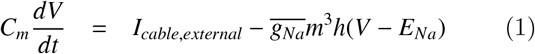

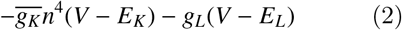

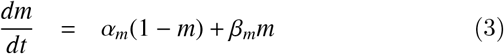

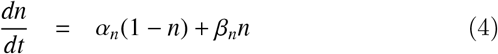

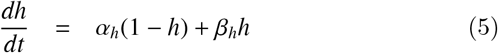

*C_m_* is the membrane capacitance. n, m, and h are dimensionless gating variables that represent the opening and closing of the gate of the ion channels. *α* and *β* are the opening and closing rates of n, m, h respectively and depending on V. The maximum conductances of sodium and potassium are 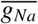 and 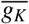, and the equilibrium potentials are *E_Na_* and *E_K_*, respectively. The conductance of leak current is expressed as *g_L_* and the equilibrium potential as *E_L_*.

For P in m, h, n (P ∈ {m,h,n}), the differential equation can be calculated in every time step as follows;

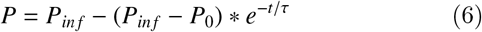

where

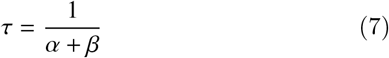

and

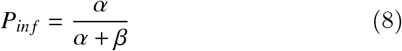

The cable equation describes propagation of membrane potential along the axis.

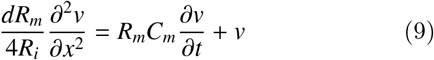

where *C_m_* and *R_m_* are the specific membrane capacitance and specific membrane resistance, *R_i_* is the specific intra-cellular resistance, *R_m_C_m_* is the time constant and d is the diameter at the position. This is a spatiotemporally continuous differential equation, which is generally difficult to solve analytically as it is. However, it is possible to obtain the solution numerically by discretizing the equation in the spatial direction. In the multi-compartment model, it is possible to obtain the solution by transforming the equation into a simultaneous equation in which the connection between compartments is represented by a sparse connection matrix.

### B. Simulator NEURON K+

The neural circuit simulator NEURON [22] is a software that supports the multi-compartment Hodgkin-Huxley type model. Although its original version was not intended for large-scale computing environments, Version 7.2 was designed to support large-scale parallelization such as MPI parallelization [23]. A simulator named NEURON K+ was developed based on NEURON 7.2, improving computational efficiency and expanding it for large-scale computing environments. A preliminary version of NEURON K+ was introduced at SC12 [15]. The current study significantly improved NEURON K+ to support the latest technology in computer architecture.

### C. Optimization of neural simulator

The excerpts from the call graph of the main functions in the simulation is as follows (Listing 1). The numbers written to the left of the functions’ name indicate the nesting level of the functions.

**Listing 1:**
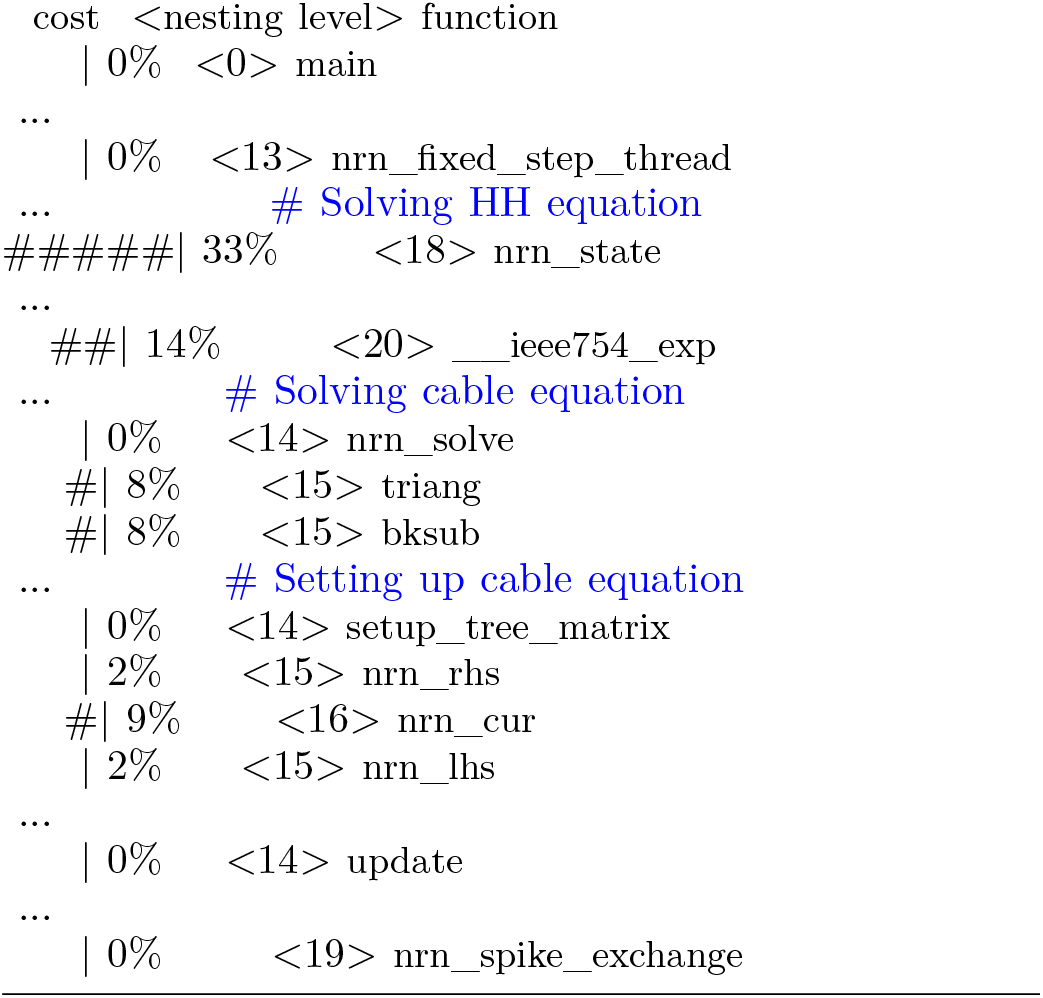
excerpts of main functions from call graphs

The main calculations in the simulation are defined by the following functions.

- nrn_rhs(): Calculation of the right-hand side of the cable equation
- nrn_cur(): Calculation of the current flowing through each compartment, called in nrn_rhs()
- nrn_lhs(): Calculation of the left hand side of the cable equation
- triang(): Calculation of the cable equation, the upper triangularization part of the Gaussian elimination method
- bksub(): Calculation of the cable equation, the part that calculates the backward substitution in the Gaussian elimination method
- nrn_state(): Calculation of the Hodgkin-Huxley equation that calculates the dynamics of the ion channel.
- nrn_spike_exchange(): Calculation of spike initiation following chemical synapse transmission between cells using MPI functions.

Source code level tuning was performed from the following perspectives. These improvements are independent of the execution environment and are effective even on generic clusters with multi-core CPUs.

#### 1) Parallelization at multiple levels

- Node and core level Hybrid parallel programming with MPI /thread-level parallelization Hybrid parallelization refers to parallelization that combines Message Passing Interface (MPI) and thread-level parallelism. Thread-level parallelism includes POSIX thread which is an API specified for POSIX systems and used by NEURON, OpenMP which is a directive based API, and other multithreading APIs.
- Instruction-level Optimization of array structure Sequential access to the memory is essential to facilitate pre-fetching to maximize the memory bandwidth. Gate parameters, m, h, and n in each compartment are stored as an “array of structure” memory layout. This data structure produces a non-sequential memory access pattern. We restructured the data structure of m, h, and n to a “structure of array” memory layout so as to enhance the compiler to SIMDize operations efficiently [15].

#### 2) Facilitation of SIMD operations

Facilitation of SIMD (single instruction, multiple data) operation was expected by bundling the variables used in the function as an array in advance. We applied this change for the functions with the highest computational cost. An example of array bundling is shown below (Listing 2 and Listing 3). In this example, the array was assigned to the pointer variable vec_d before the calculation. Because ni is an array related to compartments’ order and in regular sequence, vec_d can be accessed by systematically. This nrn_cap_jacob() function is called in nrn_lhs to compute capacitive current of each compartment.

**Listing 2:**
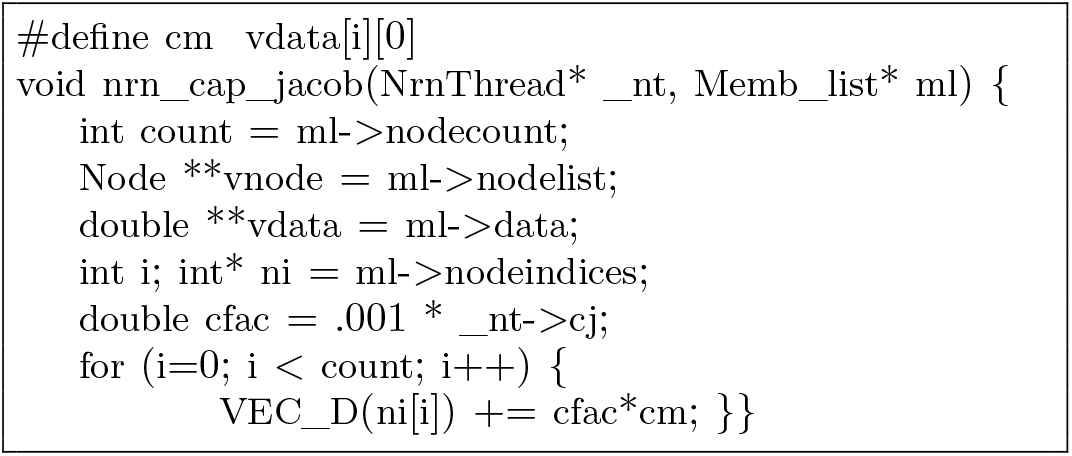
Original code

**Listing 3:**
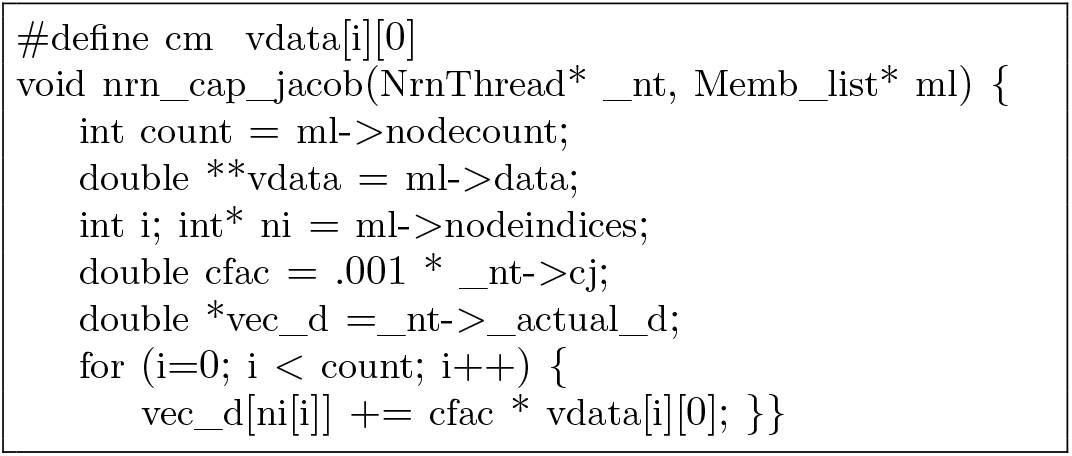
An example of array assignment

#### 3) Improvement in Memory Access Efficiency

In the functions related to the cable equation, matrices and vectors are updated in sequential calculations. The process of these functions is equivalent to setting up a simultaneous equation and solving the equation. Since the coefficient matrix is a sparse matrix resembling a tridiagonal matrix, NEURON uses a sequential update algorithm. This is because the computational complexity can be reduced to O(N) when the matrix size is N. It is predicted that the low performance of these functions is due to to irregular memory accesses during the computation as a result of matrix and vector updates being sequential operations. Referencing from child segments to parent segments occurs during the computation, but the tree structure of compartments holds a non-uniform and branching structure and is different from cell to cell, thus may result in discontinuous and irregular memory access.

In NEURON, each compartment of a cell is divided into segments with a unit called nseg, where nseg is the number of segments in the compartment, and this value must be an odd number according to the NEURON specification. Node represents the point where the channel dynamics and cable equations are calculated (Fig.2). In other words, nseg+1 operations are performed in each compartment. Therefore, as Fig.2 shows, when the node ID is 2k + 1 (where k is an integer ≥ 0), the parent node is always 2k, so that loop unrolling can be performed. “ An improvement in the function triang() is shown as an example. Listing 4 is the conventional code, and Listing 5 is the code after loop unrolling.

**Fig. 2:**
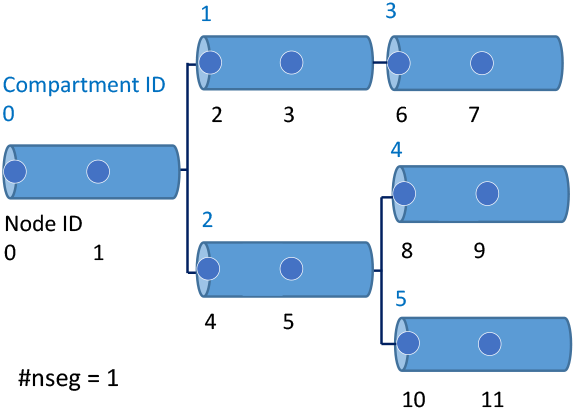
Schematic diagram of the tree structure of compartment connections

**Listing 4:**
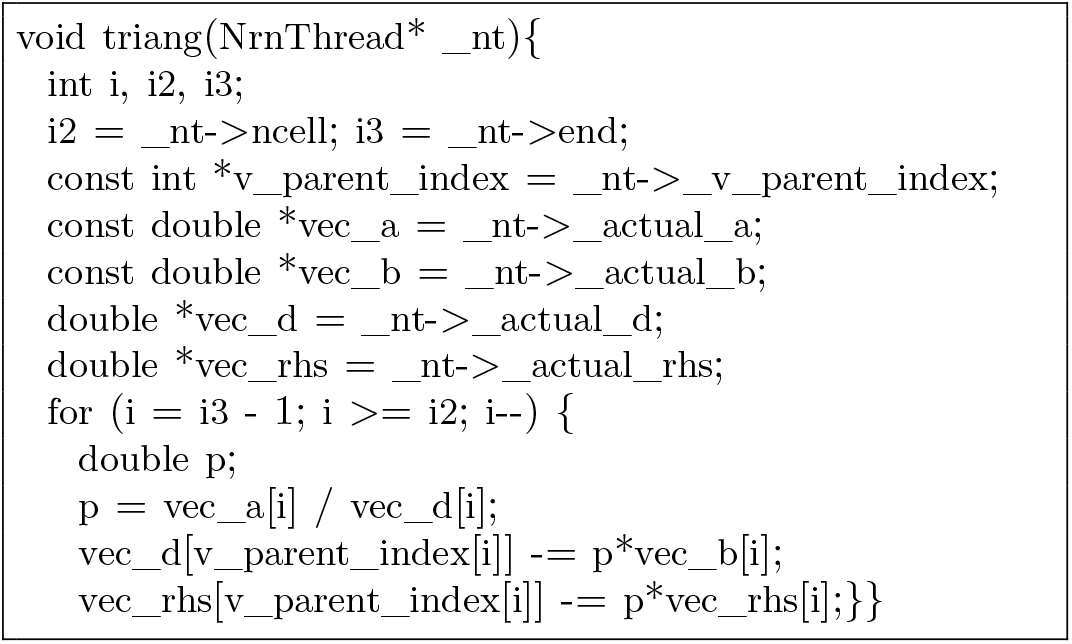
Excerpt code of triang() (original)

**Listing 5:**
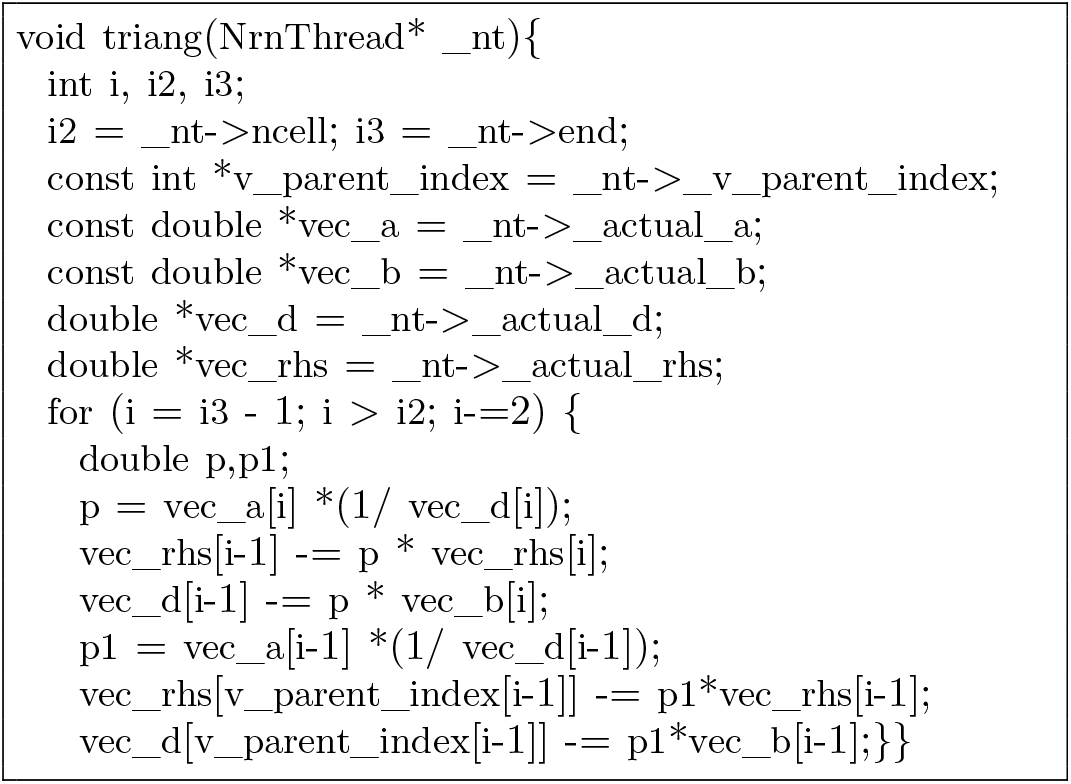
Excerpt code of triang() (modified)

#### 4) Optimization of Look-up Tables (LUTs)

Look-up tables (LUTs) were utilized in calculations of nrn_state() described in SectionIII-C. Advancing Hodgkin-Huxley equations includes operations which update m,n, and h parameters, However, the operations include other operations to update minf, mtau, hinf, htau, ninf, and ntau parameters, that involve exp(), causing relatively high computational cost as a result. Therefore, we created LUTs which corresponded to values of potential V in advance. Then we used the following equation to update the minf parameter using interpolation. We did the same for mtau, hinf, htau, ninf, and ntau parameters, also.

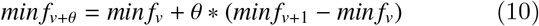

Because the arrays were indexed by potential V in a compartment and accessed by the same index, we restructured the data structure of LUTs to the “array of structure” memory layout for better access efficiency.

Conditional branching hinders vectorization of calculations and increases the stall rate of software pipelining. In the implementation of LUTs, conditional branching was used in linear interpolation. We separated this part to another loop to improve vectorization.

### D. Load Balancing

The computational cost in the simulation is roughly divided into the computations related to ion channel dynamics and cable equations for each cell and the computations related to the synaptic transmission between cells. The amount of computations required for each cell is strongly correlated to the number of compartments of the cell. In other words, when simulating a large-scale model consisting of a heterogeneous group of cells without optimization, it is expected that the cells with a large number of compartments will become will become the bottleneck factor under the normal condition that the simulation of a neuron is allocated to a core. Therefore, we used the following two methods to improve the load balance.

#### 1) Cell morphology reduction

Cell morphology reduction is a technique to reduce the number of cell compartments while taking into account their electrophysiological properties. There are two methods: branch trimming (method 1), in which a terminal branch is deleted and the volume of the deleted branch is added to its parent branch, and short-circuiting (method 2), in which an unbranched node is deleted, and the sum of the length of the original two branches becomes the new compartment length. By repeatedly applying methods 1 and 2, the number of compartments in a cell can be reduced while maintaining the general shape of the cell morphology. Fig.3 shows an example of how the number of compartments was reduced after branch trimming (method1) and short-circuiting (method2). The cell used is identified neuron BN_1056 from the silkmoth brain.

**Fig. 3:**
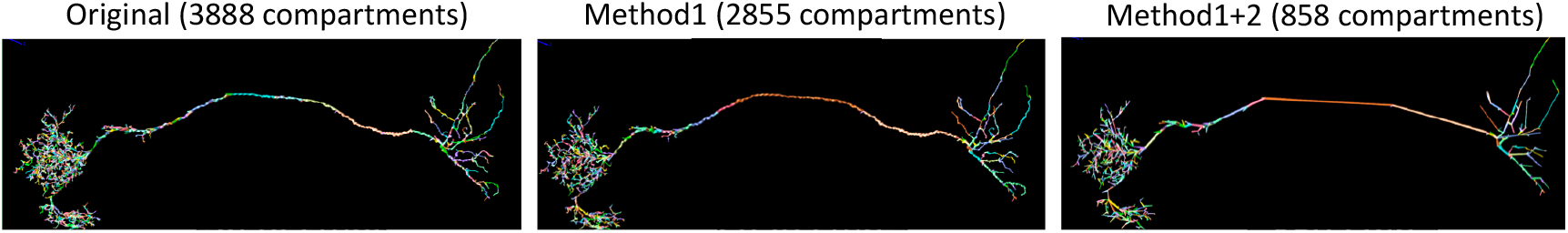
An example of cell reduction

#### 2) Cell split

Cell split is a method of dividing one cell into multiple parts and allocate the computation of each subtree to a different core [24]. We created a software scheme using the cell split technique in a large scale simulation. With this technique, sequential calculations such as cable equations can be divided among multiple cores, but if the number of subsections is too large (more than 10-20 cores), the overall execution time may increase due to synchronization-induced delays in the current implementation.

### E. Fugaku

Fugaku is a japanese supercomputer developed jointly by RIKEN and Fujitsu. A64FX [25], a many-core CPU is adopted as a processor. The features of this processor include the use of SVE (Scalable Vector Extension, with a maximum vector register size of 512), which is based on the Armv8.2-A instruction set architecture and extended for supercomputers. In addition, the SIMD width is as large as 512 bit, compared to K computer whose SIMD width was 256 bit.

- CPU
  – 48+2 assistant cores
  – 4 CMG (Core Memory Group)/CPU
    12 Cores
    HBM2 : 8 GiB
    L2 cache: 8 MiB
  – Clock frequency : 2.2 GHz
  – SIMD length : 512 bit
  – L1 Instruction cache : 64KiB /core
  – L1 Data cache : 64KiB /core
  – Arithmetic performance(Double precision) : 3.38TFLOPS
- Memory
  – Memory bandwidth : 1,024 GB/s
  – Memory capacity : 32 GiB
- Instruction set architecture: Armv8.2-A SVE 512 bit
- SVE supported vector length : 128 / 256 / 512 bit

The nodes of the Fugaku computer are connected by the Tofu interconnect D [26], which is an improved version of the Tofu interconnect installed in the K computer. Tofu-D has a 6-dimensional mesh / torus structure, which is a 6-dimensional structure in terms of hardware, but can be cut out and used in any shape as a 3D (mesh torus) space by the application.

## IV. Modeling of Neural Circuits

### A. Benchmark model

In order to evaluate the core performance of neural simulator, we constructed a homogeneous Hodgkin-Huxley circuit model as a benchmark model. The model consisted of a silkmoth neuron, BN_1056, which had 3888 compartments. The synaptic connections between multiple BN_1056s were made according to the Watts and Strogatz network [27]. The Watts and Strogatz network is a network in which one edge of the ring network is stochastically selected and its end is connected to an also stochastically selected end. This network is positioned between the ring network and the random network, and is said to resemble the neural network architecture of an actual brain(Fig.4).

**Fig. 4:**
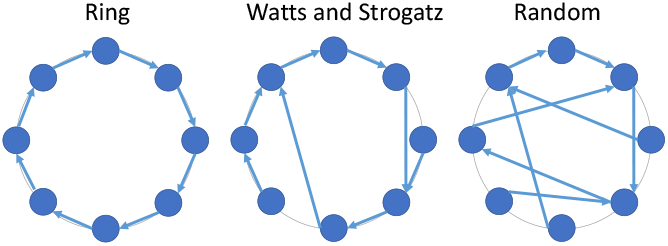
Network model

### B. Whole-brain model of Drosophila

A biologically detailed female Drosophila whole brain model for a massively parallel environment was constructed. The model was constructed in the following three steps: 1) neuron data collection and selection, 2) polarity estimation, and 3) estimation of synaptic connections.

#### 1) Neuron data

As a database of neurons, we decided to use FlyCircuit [18], a public database of Drosophila with confocal microscopic images. The major advantages of FlyCircuit are that it contains neurons from almost the entire brain and includes their neurotransmitters. Fig.5 shows an example of how information on each neuron is registered in FlyCircuit. We obtained the neuron data of female Drosophila from FlyCircuit version 1.2. 18728 cells were used as a model, after neurons with broken cell morphology that did not form tree structures and larval neurons were excluded. The data on cell morphology, neurotransmitters and innervation areas were reorganized.

**Fig. 5:**
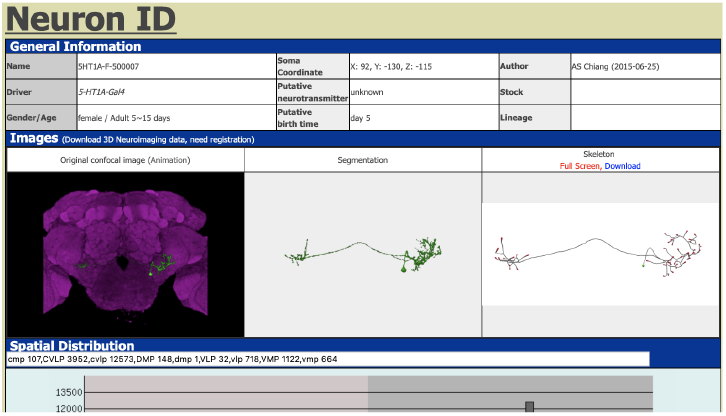
Flycircuit database (http://www.flycircuit.tw)

#### 2) Polarity identification

The polarity of each cell (axon/dendrite, etc.) was not registered in the database. Therefore, a step to estimate the polarity was necessary. As an estimation method, we used the algorithm [3], which is based on SPIN (Skeleton-based Polarity Identification for Neurons) [28]. SPIN is an open-source MATLAB software that uses machine learning to estimate cell inputs and outputs (axons/dendrites). It employs multiple regression analysis as the machine learning method applied to the morphological information of neurons such as the number of branches, and the thickness and length of branches as features. The coefficients of the multiple regression analysis were determined using Drosophila neurons with known input and output areas as training data. However, since it was known that results given by machine learning were not always correct, a manual (human) correction steps were included.

#### 3) Synaptic connection estimation

To estimate synaptic connections, we adopted Peter’s rule [29] to estimate the occurrence of synaptic contacts between neurons. This algorithm infers that the number of synapses between any given two neurons is inversely correlated to the distances between their output (axon) and input (dendrite) regions. Probability p that a synapse exists between any two compartments in output and input regions was calculated as follows.

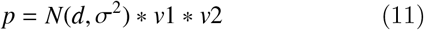

where v1 and v2 are the volumes of compartments in output and input regions, respectively, and d is the distance between the compartments. N is a normal distribution with σ = 2μ m. The normal distribution was used because the neuron data we used were mapped to a standard brain model with some error. By taking into consideration errors associated with standardization we judged that two compartments were in contact when σ = 2μm. Then we used the following criteria to select synapses for the whole-brain model.

- Threshold for the existence of synaptic connections was 3.2μm It has been reported that the upper limit of the distance at which synapses can exist between axon terminals and dendrites is approximately 1μm or less [30]. On the other hand, the local registration error in FlyCircuit’s standard brain, the database we used for this study, was 1.1μm on average, with a standard deviation of 0.2μm [18]. Since this error could not be ignored, we set an upper limit as 3.2μm for the distance at which synaptic connections existed.
- Each neuron had about 10 to 100 output synapses There are no studies conducted on the number of synapses in the whole brain of Drosophila, but there are several studies that estimated the number of synapses in particular areas of the Drosophila brain. Based on observations made on projection neurons and local interneurons of the antennal lobe (AL) using electron microscopy, the number of post-synapses ranges from several to about 100 [31]. Another study reported that the number of synapses of neurons at the calyx in the mushroom body is several dozens [32]. For insects other than Drosophila, there are studies on neurons of silkmoths. It has been reported that there are about 5 to 80 synaptic boutons for input and output neurons of the mushroom bodies [33]. The number of contacts between olfactory descending interneurons and cervical motor neurons is around 10 [34]. Accordingly, the output per neuron was set to about 10 to 100, and the number of synapses of each neuron was normalized so as to be within this range.

#### 4) Constructed whole-brain model of Drosophila

The overview of the female Drosophila whole-brain model constructed using the method above is described in this section. The model consisted of 18,728 neurons. The total number of synapses was 344,861, and the average number of synapses per neuron was 36.8. The distribution of neurons as shown in Fig.6, was fat tailed.

**Fig. 6:**
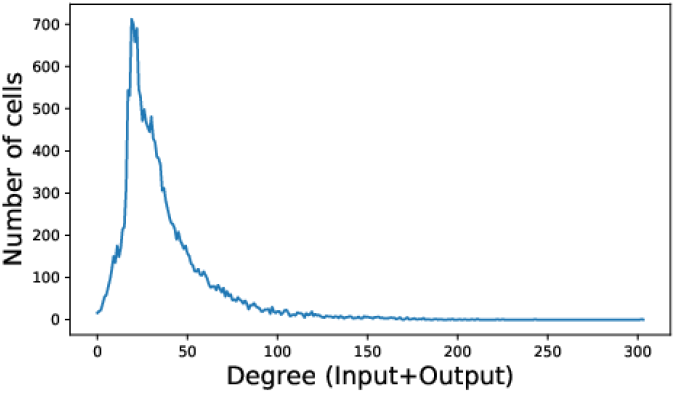
Distribution of synapse number per each neuron

We also created a brain model with cell morphology reduction and cell split as described in III-D for better load balancing. We performed the reduction process on neurons with 200 compartments or more, and repeated the operation until the number of compartments was reduced to less than 200, or the number of reductions reached 4:

- no reduction: 7445 cells
- 1 reduction: 2278 cells
- 2 reductions: 6836 cells
- 3 reductions: 1526 cells
- 4 reductions: 643 cells

The statistics for the number of compartments per cell before and after cell reduction changed as follows:

- Before cell reduction (original) Maximum value: 6180 / Mean value: 333.9 / Median: 227
- After cell reduction Maximum value: 453 / Mean value: 124.6 /Median: 126

It was seen that the overall number of compartments was reduced, and at the same time, the variability between cells was also reduced.

The histograms of the number of compartments of original model and model after the reduction are shown in Fig.7a. While the distribution of compartments in the original model is widely spread, the distribution of compartments after the reduction is concentrated, indicating that better load balancing is achieved.

**Fig. 7:**
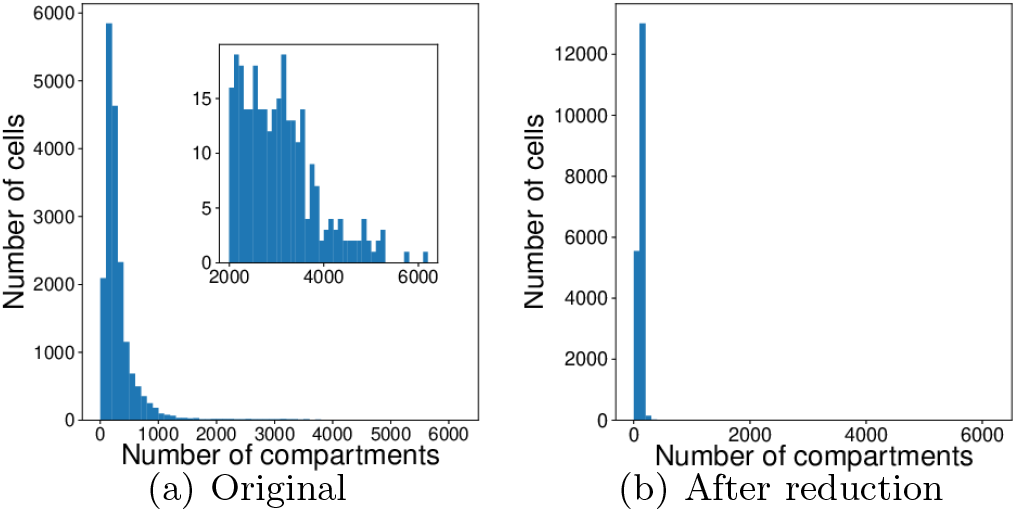
Number of compartments per cell

We also performed cell split after cell reduction on the neurons that still had more than about 300 compartments. The number of cells that were split was

- 4 divisions: 13 cells

We created the NRC (NeuRal Circuit) format which is a neural circuit description format for data description used in modeling. The format describes the cell morphology and synapse information of the cells that are assigned to each MPI process. By using this format, cell allocation to the specific core becomes easier, at the same time, modeling time can be shortened because each process reads only the necessary data. Listing 6 shows an example of a NRC file.

**Listing 6:**
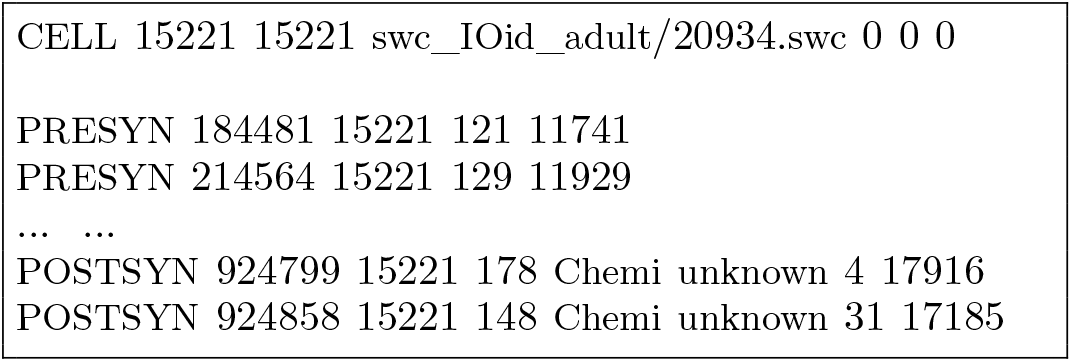
An example of NRC file

Each line represents the data of a cell as follows

- CELL: Cell definition The arguments, in order, are: cell ID (a unique continuous value assigned to each cell for management purposes, and not used in the simulation), cell gID (a unique number assigned to identify each cell in the simulator (global ID), and can be different from MPI rank ID), path to the swc file(which describes morphological information of the cell), and, coordinate position in 3D space (value of translation from the coordinates in the swc file, 0 for the purpose of this study.
- PRESYN: Presynapse definition Arguments are, in order: synapse ID, cell ID, compartment location, and postsynaptic ID (connected to POSTSYN with the same synaptic ID).
- POSTSYN: Post-synapse definition The arguments, in order, are: synapse ID, cell ID, compartment location, synapse type, neurotransmitter type, synaptic strength (not used here), and presynaptic cell ID (connected to PRESYN with the same synapse ID).

Since each NRC file corresponds to one MPI process, we prepared files for the processes and controlled the reading order to control the allocation of cells to the cores.

### C. Whole-brain simulation of Drosophila

With the whole-brain model constructed as above, we conducted a “Resting State” simulation. Resting State refers to a state in which there is no external input such as sensory input, and is the basic state for the brain. This was achieved by implementing the input to each cell as follows.

- Stimulus: Gaussian white noise current We applied a current injection that follows a constant Gaussian distribution with mean zero and standard deviation greater than zero.
- Stimulus point: 1 compartment/cell One compartment from the center part for local interneurons, and one compartment from the endpoint of a dendrite for other neuron types, were used as input targets.

## V. Results and Discussion^1^

### A. Improvement of neural simulator

#### 1) Source code level tuning

The results of optimizing the NEURON K+ code at the source code level are shown in this section. The simulation of a benchmark model with one cell was run by one core in Flat MPI. The simulation conditions were one cell with 10 synapses of the benchmark model and a simulation time of 1 second. Then, we ran NEURON 7.2 under the same conditions for comparison. The results were given by Fugaku’s profiler. The measurement range was the entire simulation excluding the time for modeling and initialization.

The improvements in the simulation is shown in Fig.8. The execution time was reduced to approximately 13% compared to NEURON 7.2.

**Fig. 8:**
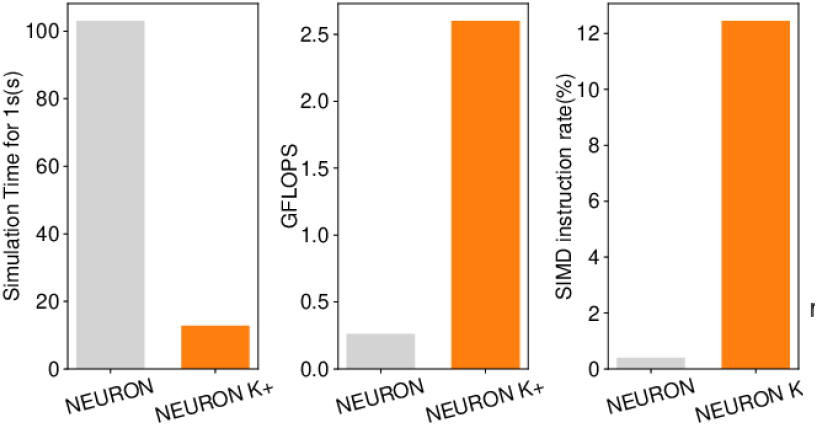
Improvement of simulator

We selected functions that were highly responsible for computational cost without optimization, and measured, for each selected function, the time required for completion, before and after the optimization. As shown in Fig.9 and Fig.10, both FLOPS and SIMD operation rate were improved. The majority of the computational cost was due to exponential functions included in nrn_state(). Our counting from the pseudo-code shows the BF ratio on one step of this function that means the calculation of Hodgkin-Huxley model, is 0.903 (130 floating-point operations / 144 byte when one exponential function is evaluated as 30 operations) but the measured value from the profile from the benchmark simulation is 1.8 (4.68 GFLOPS/8.50 GByte at 48 core execution). However, the measured value from the profile from the benchmark simulation is 1.8 (4.68GFLOPS/8.50GByte at 48 core execution). Our performance tuning with large SIMDization improved the performance of nrn_state() and nrn_rhs() to approach the main memory bandwidth limitation of Fugaku (Fig.12). The original NEURON code uses a breadth-first search algorithm for detecting child nodes. We tried replacing this by a depth-first search for the effective memory access but significant improvement of the performance were not observed in our benchmark simulations. We also tried to replace the triang() and bksub() functions that solve the cable equation by parallelized algorithm by using C-SSL2, which is a mathematical library, but the performance were lower than our case. The performance comparison with multiple morphology reduction form out benchmark neuron show that simulation performance little depend the complexity but compartment number. Especially for neurons that have the large number of compartments the performances were proportional to the compartment number (Fig.11).

**Fig. 9:**
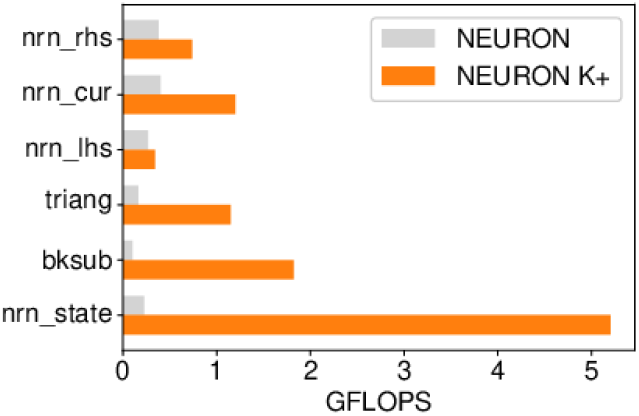
FLOPS for each function

**Fig. 10:**
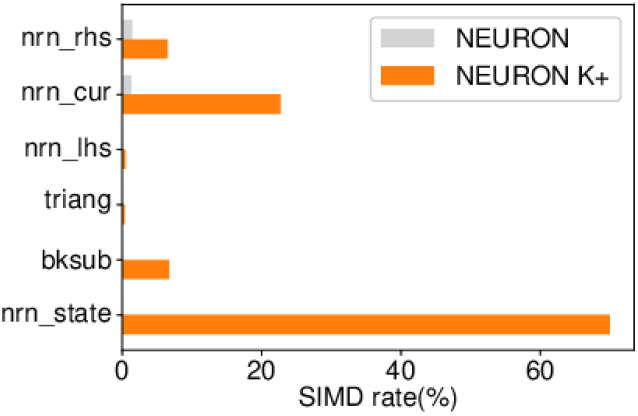
SIMD operation rate for each function

**Fig. 11:**
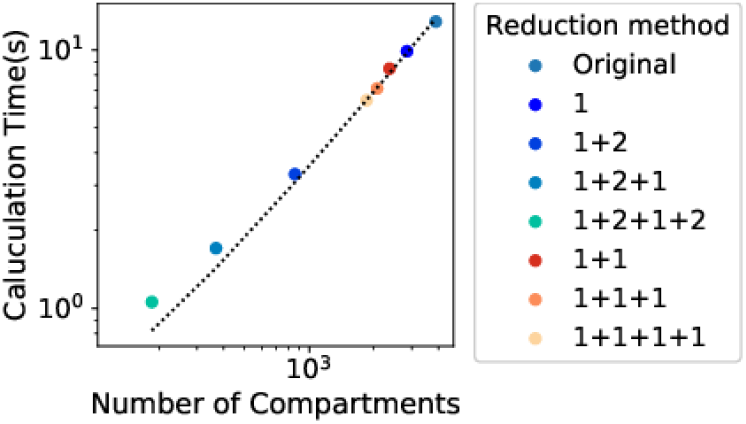
Relationship between calculation time and number of compartments

**Fig. 12:**
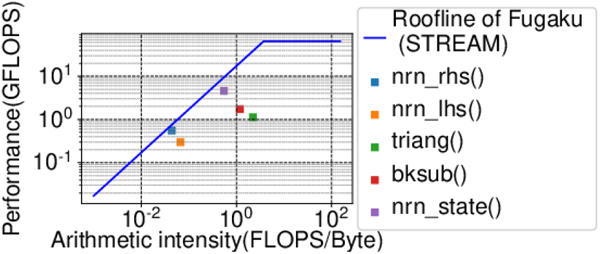
Roofline analysis

#### 2) Weak scaling

We performed a large weak scaling experiment on Flat MPI. The simulation time was 1 second.

Although the execution time increases as the number of cells increased as shown in Fig.13, the parallel efficiency was high at 0.999998 with 480,000 cores. Fig.14 shows total FLOPS of the simulation in each number of cells. We also achieved above 630 TFLOPS with 480,000 cores. Next we conducted weak scaling experiments with longer time steps, dt=0.25. Under this condition, calculation time was reduced to about 1/6 to 1/3 compared to time steps of dt=0.025. We tried hybrid MPI with 6 threads for 1 cell, However, using hybrid MPI with 6 threads per cell lead to significantly longer simulations times in particular with a large cell numbers (>50000). Fig.15 shows the components of the simulation time. “Other” part of the simulation includes computation of each cells such as calculation of Hodgkin-Huxley equation and cable equation. When the number of cells was large, the total computational time was increased because MPI functions used in spike_exchange() took larger cost.

**Fig. 13:**
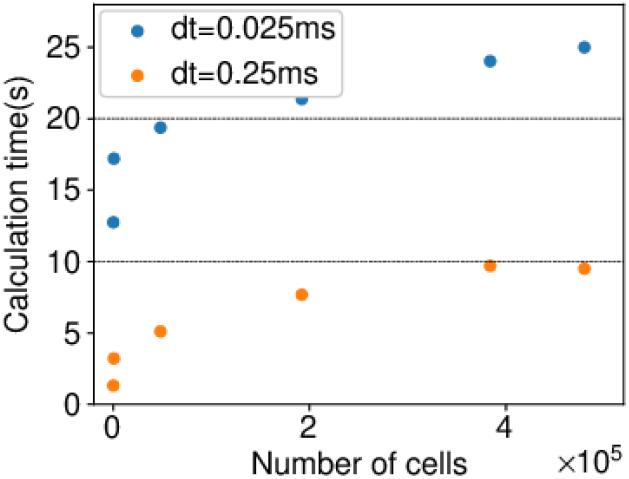
Calculation time in weak scaling

**Fig. 14:**
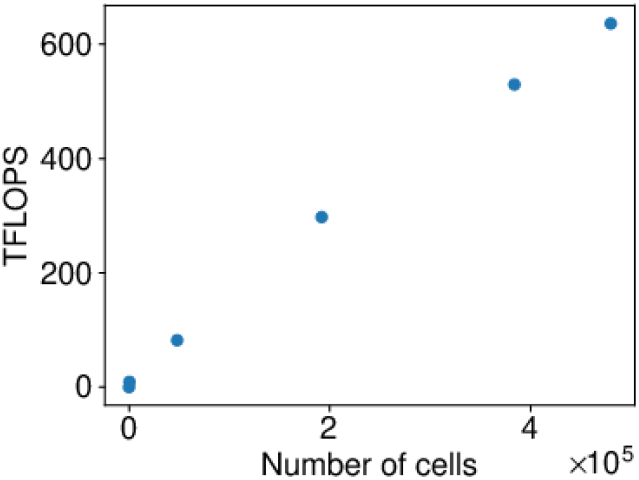
FLOPS in weak scaling

**Fig. 15:**
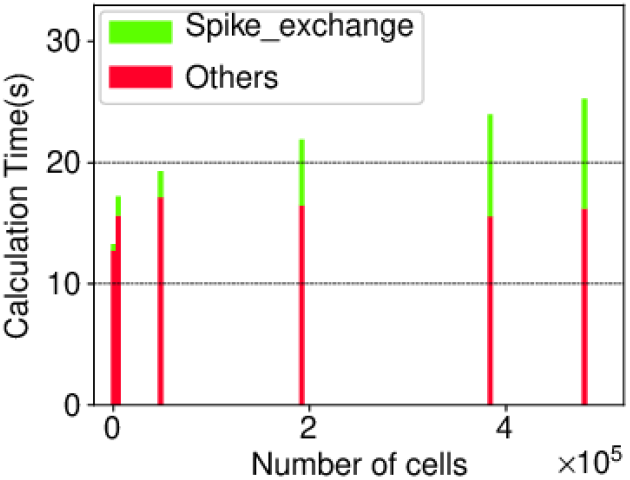
Computation by components in weak scaling

#### 3) Strong scaling

Strong scaling measurements were conducted with a benchmark model. The simulation model was the benchmark model with 20,480 cells which was fixed throughout the experiment, and we changed the number of cores. Experimental results are shown in Fig.16. We conducted experiments under four different conditions: cells without split with Flat MPI (indicated as “Cell without split” in the figure), cells without cell split with Hybrid MPI (indicated as “Without cell split + Hybrid MPI” in the figure), cell split with Flat MPI (indicated as “Cell split” in the figure), and cell split with hybrid MPI (indicated as “Cell split + Hybrid MPI” in the figure). “Cell without split” resulted in good scaling. Under this condition, however, it is needless to say that the maximum number of cores used was limited to the number of cells. On the other hand, cell split enabled the simulator to use cores that are up to 12, because one CMG contained 12 cores as mentioned in Section.III-E. Hybrid MPI enabled the simulator to operate multiple threads in one core. The results showed that “Cell split” performed higher in efficiency than “Hybrid MPI”. For “Hybrid MPI”, 8 threads took more calculation time than 6 threads and 12 threads, and one possible reason for this was that some cells were allocated across different CMGs, causing additional delays. On the other hand, the results of “Cell split”, shows that 12 splits of cell took longer calculation than 8 splits. In general, as the number of cell split increases, the time required for the synchronous process increases, and it is considered that this was the case for the time increased. As for the “Cell split + Hybrid MPI”, where we used 12 cores, 4 splits (4 MPI) with 3 threads in parallel, we achieved a parallel efficiency of 0.79.

**Fig. 16:**
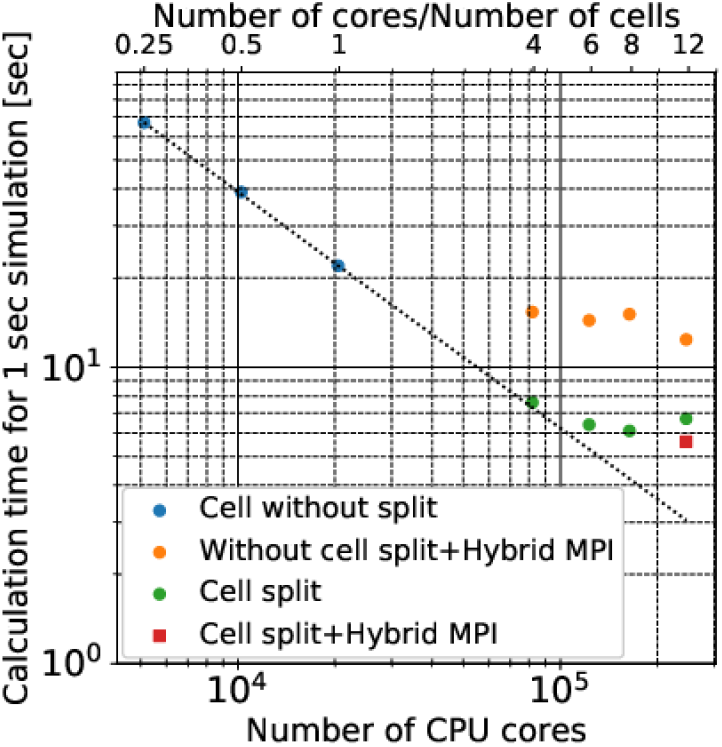
Strong scaling experiment with 20,480 cell Upper x-axis indicates that the number of cores used is n times the number of cells.

### B. Whole-brain simulation of Drosophila

We simulated the Resting State condition using the constructed whole-brain model of female Drosophila with the optimized neural simulator. Fig.17 shows the visualization of potential V of each neuron cells during the simulation. With a step time, or a discretization interval, dt= 0.025ms, the time required to run the simulation for 1 second of brain activity was 21.3s for the original model without cell reduction and cell split, and the execution time was 2.63 s for the model with cell reduction and cell split (Fig.18).When step time dt was increased to 0.25ms, the simulation execution time for 1 second was 0.97 seconds in the model, and real-time simulation was achieved. MPI_Send/Receive was faster than MPI_Allgather as the type of MPI communicator function in all three cases. This indicates that point-to-point communication in conjunction with the interconnect of Fugaku was better suited to simulate insect brains with sparser synaptic connectivity compared to vertebrate brains. The modeling time was also reduced to about 6 seconds in this case using the NRC format. The average spike firing rate in this Resting State simulation was 3.2±8.6 Hz. The spontaneous firing in Drosophila has been reported in several studies. For the projection neurons of the antennal lobe (AL), the mean rate of firing in the absence of stimulation was reported to be 4 ± 0.7 Hz [35]. For the local interneurons (LNs) of the AL, spontaneous firing was reported to be 4.0 16.8 Hz [36].

**Fig. 17:**
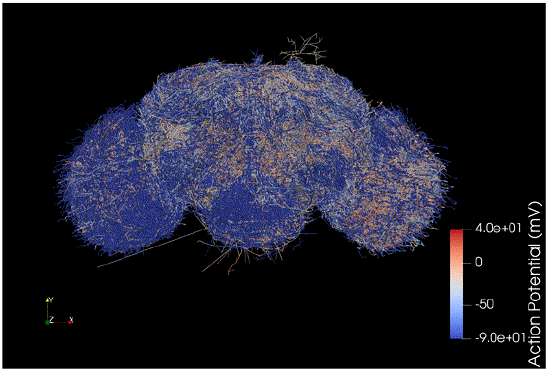
Visualization of the simulation

**Fig. 18:**
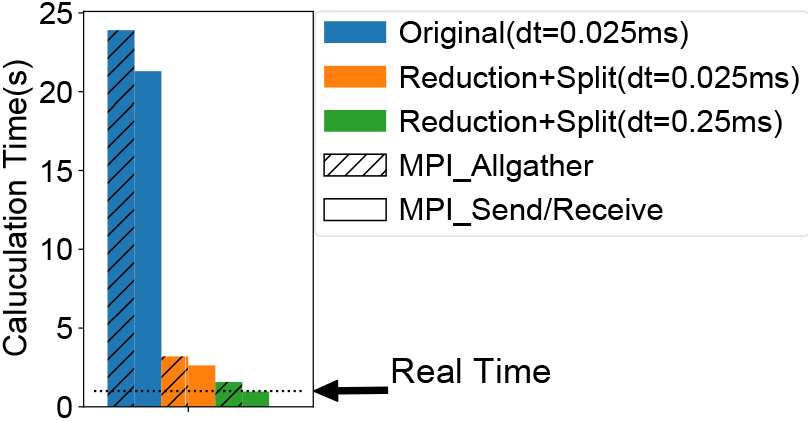
Simulation time of Resting State

Our Resting State simulation thus reproduces the overall behavior of resting state neurons in the Drosophila brain.

### C. Olfactory system

Olfaction is an essential sensory modality for insects. They approach or avoid plants, other animals, and objects by identifying odors with innate or learned connotations. The proboscis extension reflex (PER) where odors act as conditioned stimulus (CS) and taste can act as unconditioned stimulus (US), is a typical and well studied behavior as an elemental learning systems [7] in many insect specie. We especially reconstructed the neural circuit of the olfactory pathway, from receptor neurons through the antennal lobe to the mushroom body, the higher center in insect brain that is related the odor-taste associative learning (Fig.19). Required morphology of each neuron (uni-glomerlar projection neruons; uPNs, multi-glomerlar projection neruons; mPNs, local interneurons; LNs, Kenyon cells; KCs, dorsal paired medial neurons; DPMs, dopamine neurons; DANs, mushroom body output neurons; MBONs) were mainly obtained from FlyCircit though some part of these were downloaded from FlyEM and translated to the one under the coordinates system of FlyCircuit. ORNs were built as simple rods that enter each glomerulus in the antennal lobe. The olfactory responses in ORNs were generated using Poisson processes to reproduce time courses reported in [37]. Physiological properties of PNs and LNs were constructed by adjusting maximal conductances of Na^+^, K^+^, Ca^2+^, and AHP ion channels heuristically as [38] and [39]. The other properties were build as described above. Reward signals were applied to DAN. A standard STDP (Spike-timing dependent synaptic plasticity) mechanism was implemented in the MB-lobes. In this simulation we succeeded to detect olfactory responses that were reinforced by reward signals in MBON output neuron. Synaptic weights between KC and MBON6 were increased during training(Fig.20), and we confirmed that the acquired a strong response to the rewarded olfactory input after training(Fig.21).

**Fig. 19:**
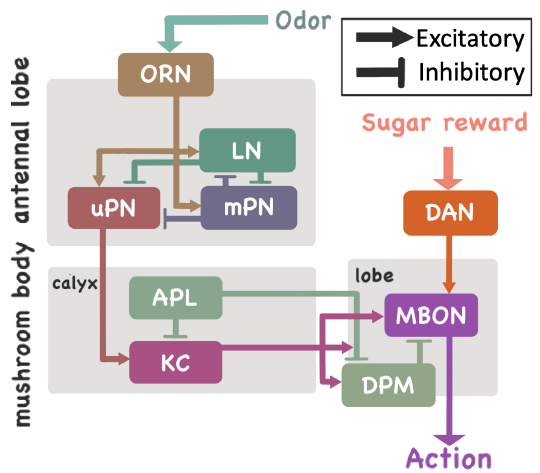
The neural network involved in odor-taste associative learning

**Fig. 20:**
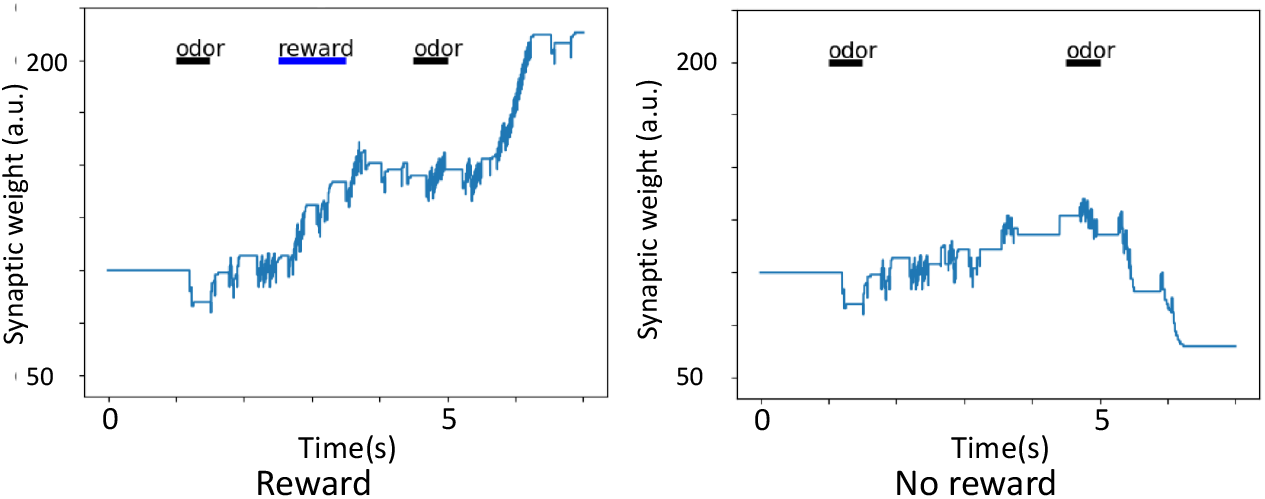
Changes of synaptic weights with and without reward input

**Fig. 21:**
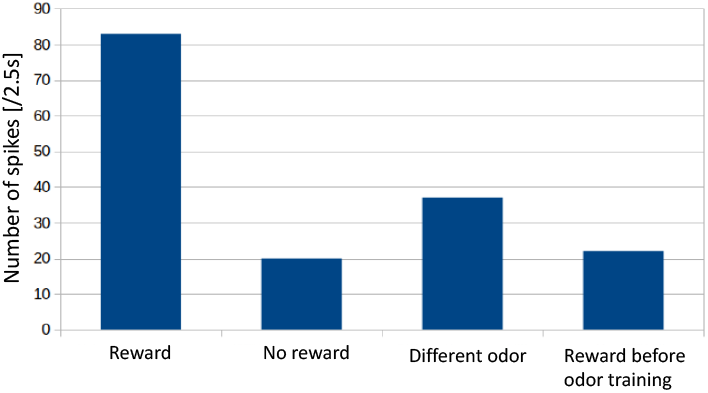
MBON activities in relation to olfactory stimuli and reward

## VI. Conclusions and Future work

In this study, based on NEURON, the most popular neuron/neural circuit simulator supporting multi-compartment Hodgkin-Huxley models, we developed NEURON K+, that is optimized for massively parallelized neural circuit simulations and implemented a simulation of an insect brain.

As a result, we have achieved a high performance of 630 TFLOPS and the parallel-computing efficiency of 0.99998, with 480,000 cores contained in 10,000 nodes, in a weak scaling experiment with a benchmark model in Fugaku. This is equivalent to an improvement of 8.6 times in execution speed and 10 times in calculation efficiency compared of the constructed a female Drosophila’s whole-brain model to NEURON 7.2. Introducing the cell division technique in the neural circuit simulations realized real-time or near real-time simulation. Further improvements in the neural circuit simulator are expected in future work by addressing the following challenges. We have shown that calculation speed of ion channel kinetics and part of the cable equation are currently limited by main memory bandwidth and that smart and efficient use of the L2 cache for membrane states can improve the calculation calculation speed for the simulation of these elements of neural models. Since nodes of the Fugaku computer are in torus structure, communication time is expected to be reduced through the optimization of cell placement with respect to the core. It is also considered necessary to devise ways to speed up communication within the same CMG. This large-scale model is a detailed model that incorporates cell polarity and synaptic connections supporting large-scale parallel computing environments. It is possible to build similar models for large-scale computational environments for other organisms as well, given that there are cell morphology data, and polarity data or training data for estimation. Of cause this complex and nonlinear simulation include some error in many biophysical parameters, for example ion channels’ spatial distributions, and these kinetics, synaptic weights of each synapse that we heuristically determined. However, odor-taste association learning was successfully performed by our simulation showing that the bottom-up reconstruction of insect brains by the biophysically detailed model based on optical microscopy morphology data of brain neurons is a fairly feasible and have possibility that can reproduce brains’ functions. Because the whole brain simulation can be achieved in a smaller scale in the whole Fugaku, parameter estimation using evolutionary algorithm can be possible. We developed parameter estimator based on an evolutionary algorithm CMA-ES [40] [41]. In Fugaku that allows dynamical and hierarchical structure of MPI, we will be able to develop the parameter estimator of whole insect brain simulation easily. Moreover simultaneous executions of real time simulation will allow real time data assimilation for individual real neuron/brain simulations [42] [43] using physiological measurements. As a first step towards the simulation of brain functions of Drosophila’s whole brain, the results we have obtained are overall satisfactory.

## Footnotes

1 The results obtained on the evaluation environment in the trial phase do not guarantee the performance, power and other attributes of the supercomputer Fugaku at the start of its public use operation.

